# Technique for rapidly forming networks of microvessel-like structures

**DOI:** 10.1101/2023.06.22.546165

**Authors:** Sarah A. Hewes, Fariha N. Ahmad, Jennifer P. Connell, K. Jane Grande-Allen

## Abstract

Modelling organ-blood barriers through the inclusion of microvessel networks within *in vitro* tissue models could lead to more physiologically accurate results, especially since organ-blood barriers are crucial to the normal function, drug transport, and disease states of vascularized organs. Microvessel networks are difficult to form, since they push the practical limit of most fabrication methods, and it is difficult to coax vascular cells to self-assemble into structures larger than capillaries. Here we present a method for rapidly forming networks of microvessel-like structures using sacrificial, alginate structures. Specifically, we encapsulated endothelial cells within short alginate threads, then embedded them in collagen gel. Following enzymatic degradation of the alginate, the collagen gel contained a network of hollow channels seeded with cells, all surrounding a perfusable central channel. This method uses a 3D printed coaxial extruder and syringe pumps to generate short threads in a way that is repeatable and easily transferrable to other labs. The cell-laden, sacrificial alginate threads can be frozen after fabrication and thawed before embedding without significant loss of cell viability. The ability to freeze the threads enables future scale up and ease of use. Within millifluidic devices that restrict access to media, the threads enhance cell survival under static conditions. These results indicate the potential for use of this method in a range of tissue engineering applications.

**Impact Statement:** Generating microvascular networks is a challenge in tissue engineering. Popular 3D bioprinting techniques use sacrificial structures to generate vascular networks, but those approaches can require expensive equipment and extensive troubleshooting. In this article, a method is presented for easy and rapid generation of centimeter-scale microvascular networks using endothelial cells frozen within sacrificial structures. The method for fabricating and freezing the sacrificial threads of alginate containing cells using a 3D printed coaxial extruder is also described. Experimental results indicate that these microstructures have the potential to enhance cell survival in environments with restricted access to media.

## Introduction

Blood vessel networks are hierarchical; larger blood vessels branch into microvessels that branch into capillaries. Numerous *in vitro* and *in vivo* studies have shown that endothelial cells will self-assemble to form a capillary network within a scaffold material, such as collagen, even over centimeter scales.^1^ These self-assembled capillaries form with diameters of around 5-30 µm.^2^ Large vessels can be created by mechanical means, such as molding or 3D printing.^3,4^ However, in extrusion bioprinting, the diameter of the nozzle used to print live cells is typically limited to a minimum size around 100 µm^5^ due to mechanical forces that negatively impact cell viability at smaller diameters.^6^ Microvessel networks containing blood vessels larger than capillaries are the most difficult to form since they push the practical limit of most fabrication methods and since it can be challenging to coax cells into self-assembling vascular networks at this scale.

One approach to establish 3D vasculature *in vitro* is to fabricate hollow spaces in a hydrogel where blood vessels are desired. Generally, endothelial cells cultured *in vitro* tend to spread to cover available space before branching out through a solid matrix. Many investigators take advantage of this behavior by seeding endothelial cells within pre-formed vascular structures.^7–9^ Sacrificial materials are typically employed to create these spaces within the bulk material; after the sacrificial material solidifies, it is subsequently dissolved by chemical, thermal, or enzymatic means. To seed cells within a structure prepared with sacrificial material, the channels must be able to be perfused. In a complex structure with disconnected forms, direct infusion of cells becomes impractical, so the cells are sometimes encapsulated within the sacrificial structures.

Alginate is a common choice for a sacrificial material because the conditions for its gelation and degradation are compatible with live human cells. Endothelial cells encapsulated within or grown on the outside of alginate beads or fibers have been used to generate vasculature *in vitro* in combination with gelatin, fibrin, or collagen.^10–14^ Though cells cannot degrade alginate and thus cannot proliferate within alginate structures, alginate is a highly biocompatible material that is widely used in tissue engineering.^15^ Despite its limited utility for cell scaffolding, alginate is an excellent material for temporary cell encapsulation, due to its reversible gelation properties. Although a calcium chelator such as sodium citrate is a popular method for reversing alginate gelation, enzymatic degradation using alginate lyase (alginase) is an alternative approach to assure that the alginate structure does not re-form when calcium-containing media is introduced to the system. Alginate lyase has not been found to negatively affect the viability, proliferation, or sprouting potential of endothelial cells^16^.

The goal of this work was to create a platform for studying vascularized tissues *in vitro*. Here, we describe a method for rapidly forming microvessel networks from cellularized, sacrificial structures that can be premade, stored, and used from frozen aliquots.

## Method

### Fabrication of coaxial extruder

A coaxial extruder was designed to produce alginate structures of a desired diameter and length (**Figure 1A**). The simple extruder consisted of a needle positioned lengthwise through the center of a larger tube. A mixture of cells and alginate (Sigma Aldrich, St. Louis, MO) was flowed through the needle while calcium chloride was flowed through the larger outer tube, creating a sheath of CaCl_2_ around an alginate core. The alginate core was quickly crosslinked by the calcium ions to form a thread-like, alginate structure encapsulating cells. The minimum goal for the extruder design was to be able to produce at least 1 mL of threads in under 30 min, while using at most 120 mL of CaCl_2_. To prevent shear-induced cell death, the needle diameter had to be greater than 100 μm (32 gauge). To achieve alginate threads with diameters smaller than the needle diameter of 432 µm (23 gauge), the flow rate within the CaCl_2_ sheath needed to be substantially higher than the flow rate in the alginate core.

**Figure 1.**
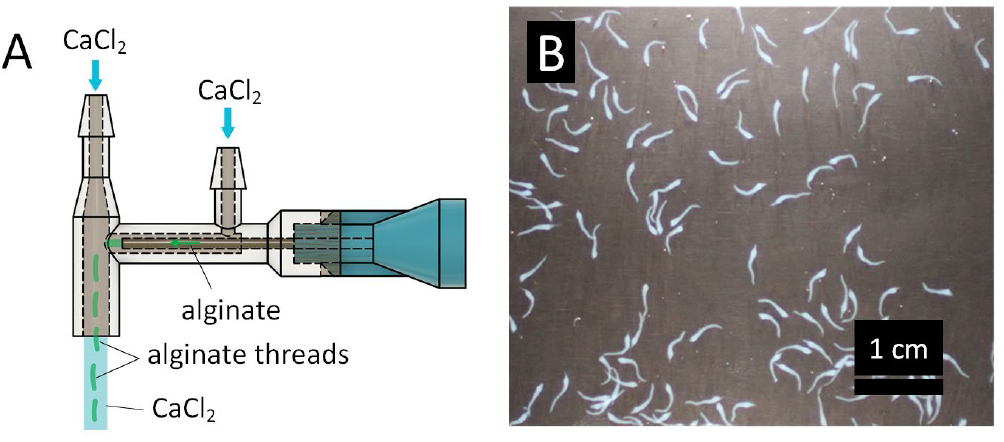
Coaxial extruder and alginate threads. A) Schematic of coaxial extruder used to make alginate threads. B) Macroscopic image of alginate threads.

The extruder was also designed to be easily reproducible. The extruder sheath was printed with a Form 2 SLA printer with Dental SG resin (Formlabs, Boston, MA), thoroughly washed with isopropyl alcohol, cured, and attached to a 23 gauge ½” length blunt tip needle (McMaster-Carr, Elmhurst, IL) using Loctite GO2 glue (Loctite, Westlake, OH). The use of a commercial SLA printer, together with the alignment features on the extruder sheath, allowed for consistent reproduction of these core-sheath extruders.

Once prototypes of the simple coaxial extruder could produce continuous alginate threads of consistent diameter, the crossflow portion of the extruder was added. The crossflow periodically dragged the threads downstream away from the needle abruptly, thereby shearing them into threads of consistent length (**Figure 1B**). The CaCl_2_ port nearest to the Luer-lock connector of the needle created the fluid sheath that formed the threads. The further CaCl_2_ port created the crossflow to shear the threads off at regular lengths. The tip of the needle was positioned 2 mm from the center of the crossflow and the crossflow channel had a 1 mm radius. Extruders with crossflow center to needle tip distances of 0 – 6 mm were tested. The length of the alginate threads appears to scale with the distance from the extruder, except that very small diameter threads were produced when the tip of the needle was at the edge of the crossflow. The distance between the tip of the needle and the crossflow portion was found empirically, with the goal of producing threads around 4 mm long that would be less likely to tangle and bind together, but long enough to produce microvessels resembling a physiological network.

### Fabrication of alginate threads

Alginate threads were fabricated using the 3D printed coaxial extruder. After assembling the extruder and allowing the glue to cure for 24 hours, the extruder assembly and all tubing was autoclaved. To encapsulate HUVECs within alginate threads and facilitate the formation of threads significantly smaller than the needle diameter, the alginate flow rate was set at 30 mL/hr and the CaCl_2_ flow rate was set at 40 mL/min at each inlet port. Since the sheath flow rate was significantly higher than the core flow rate, two syringe pumps were used. One syringe pump dispensed the alginate and cell mixture to the needle with a 5 mL syringe, and the other dispensed CaCl_2_ from two 60 mL syringes that were connected to the two extruder ports (**Figure 1A**). A 2% solution of alginate buffered with 15 mM HEPES containing green fluorescent protein-expressing human umbilical vein endothelial cells (GFP-HUVECS) (Angio-Proteomie, Boston, MA) at a concentration of 3 million cells/mL was flowed through the needle. A solution of 80 mM CaCl_2_ in 15 mM HEPES, adjusted to pH ∼7, was flowed through the other two ports. Cell-laden alginate threads were collected in conical tubes. After allowing the threads to settle, the calcium chloride was aspirated. The threads were then collected in cryovials and resuspended in 10% DMSO, 90% FBS. The threads were frozen for 24 hours at -80°C and then stored in liquid nitrogen for up to 7 months.

### Design of vascular network culture device

A flow device to support the collagen-embedded vascular network was created with several considerations in mind. To ensure that device would sustain a perfusable network for the duration of the cell culture experiments, it was designed to include a media reservoir and to have the capacity for recirculating flow at the desired flow rates (**Figure 2A-C**). The surface area of the device was approximately 1.28 cm^2^ (four times the area of one well of a 96-well plate) to accommodate a volume of cells that would enable the collection of abundant mRNA and proteins for biomarker detection assays. A central channel was included along the length of the device, designed to be 500 µm, which was considered sufficiently large to avoid clogging with bubbles or cell clumps, yet small enough to avoid collapse when fabricated from rat tail collagen (Cat # 354249, Corning, Glendale, Arizona). The height of the collagen gel was limited to 2 mm due to imaging constraints.

**Figure 2.**
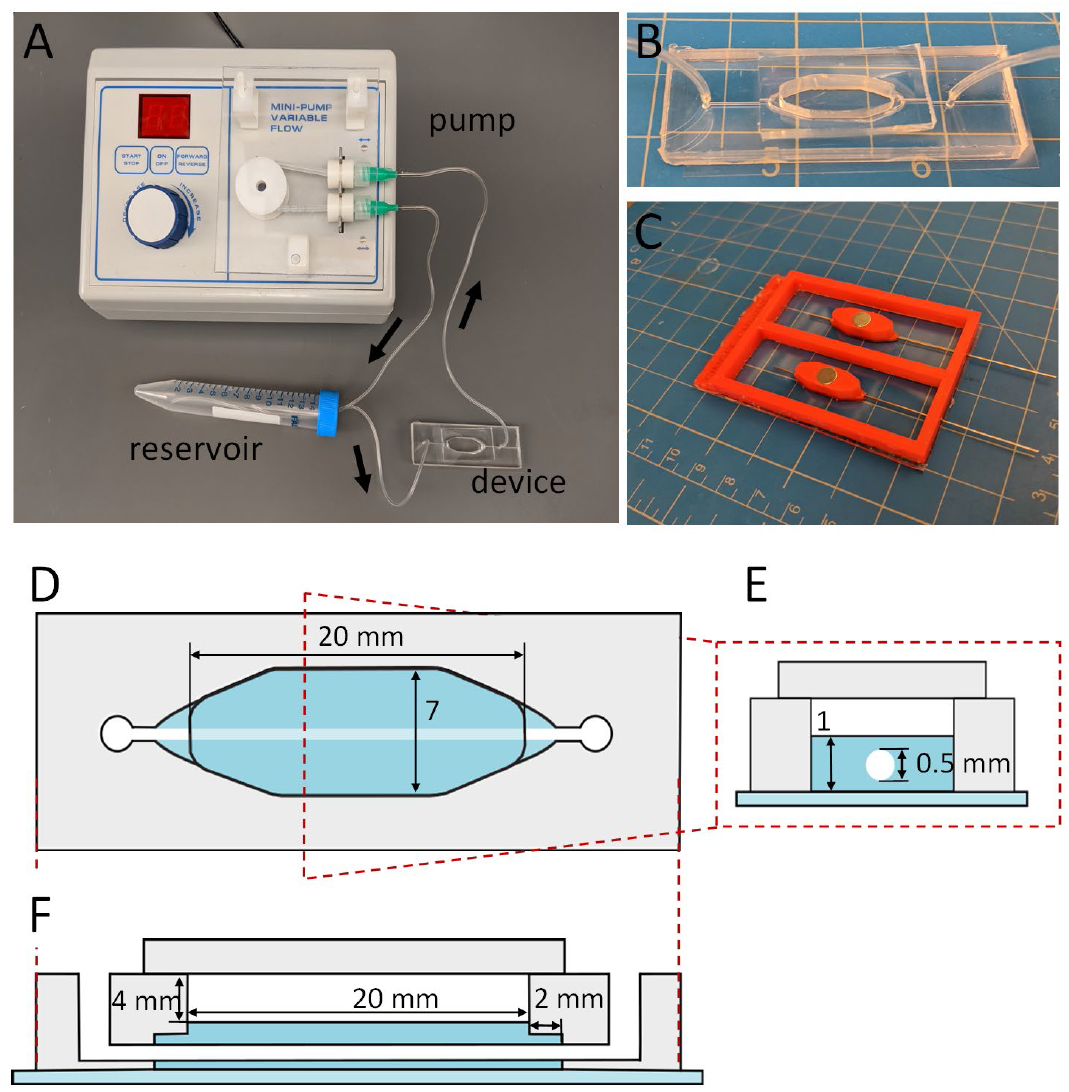
Schematics of flow device and flow loop. A) Flow loop assembly labeled with arrows indicating flow direction. B) Close up of millifluidic device (grid is 0.5 inch squares). C) 3D printed mold for millifluidic device. D) Top-down schematic showing dimensions of millifluidic device with red dashed line indicating cross section in Panels E and F. Gray indicates PDMS, blue indicates collagen, and light blue indicates glass. E) Cross section of the center of the millifluidic device in D. F) Longitudinal section, perpendicular to the dashed red line in D.

To fabricate the device, 3D printed negative molds were printed using polylactic acid (PLA) and attached to glass with Loctite GO2 glue (**Figure 2D**). After allowing the glue to set for 24 hours, polydimethylsiloxane (PDMS) (Dow, Midland, MI) was mixed in a 1:10 ratio of crosslinker to prepolymer, bubbles were removed using a vacuum chamber, and the PDMS was poured into the molds with a 508 µm diameter wire (McMaster-Carr) inserted (**Figure 2D)** to create a channel for the needle to be inserted in a later step. After removing any remaining bubbles, the molds were covered with a plastic sheet to create a flat top surface. The PDMS was cured at room temperature for 48 hours. After removing the PDMS from the negative molds, the PDMS was baked for 1 hour at 90°C. A 1.5 mm biopsy punch was used to create ports on either side of the chamber, connected to the central channel. The PDMS device was then plasma bonded to 24 mm x 60 mm, No. 2 cover glass (VWR, Radnor, PA) using a laboratory corona treater (Electro-Technic Products, Chicago, IL). A thin, No. 2 cover glass was used to facilitate imaging. Small slabs of PDMS, used for sealing the device chamber, were cut from cured PDMS in 4 mm thick rectangular slabs approximately 15 mm by 30 mm. The devices were autoclaved before use.

### Culturing of HUVECS

GFP-HUVECs were cultured in EGM-2 (Lonza, Basel, Switzerland) media on uncoated tissue culture flasks. Media was changed every second day. After a total of three days of culture in an incubator (37°C, 5% CO_2_) or upon reaching 80% confluence, the cells were passaged with a 1:6 split. GFP-HUVECs were not allowed to become confluent. For passaging, cells were detached from the flask using 0.05% trypsin, which was then neutralized with EGM-2. The cells were used in experiments at passage seven.

### Viability studies

The ability to retrieve cell-containing sacrificial structures from frozen aliquots greatly simplifies experiments. To verify that the cells were not compromised by this process, cell viability was assessed using Calcein-AM (Invitrogen, Waltham, MA) and Ethidium Homodimer-1 (Eth-D) (ThermoFisher, Waltham, MA). Cells that had been frozen in liquid nitrogen for a minimum of 24 hours were thawed quickly in a 37°C bead bath. Eth-D and Calcein-AM were added in concentrations of 2 µl/mL and 1 µl/mL, respectively, and incubated with the cells for 10 minutes before imaging on a fluorescence microscope. Cells were counted using CellProfiler (Broad Institute, Boston, MA).^17^ Cell viability was evaluated for cells after passaging, cells that had been thawed after freezing, cells that had been encapsulated, and encapsulated cells that had been thawed after freezing. The results were collected over three separate encapsulation experiments.

### Embedding in collagen

To evaluate whether the encapsulated structures would promote the formation of a microvascular network, cells were seeded under two conditions. GFP-HUVECs were randomly seeded within a collagen gel, or a similar number of GFP-HUVECs that had been encapsulated within sacrificial alginate threads were mixed within the collagen gel as described below (**Figure 3**).

**Figure 3.**
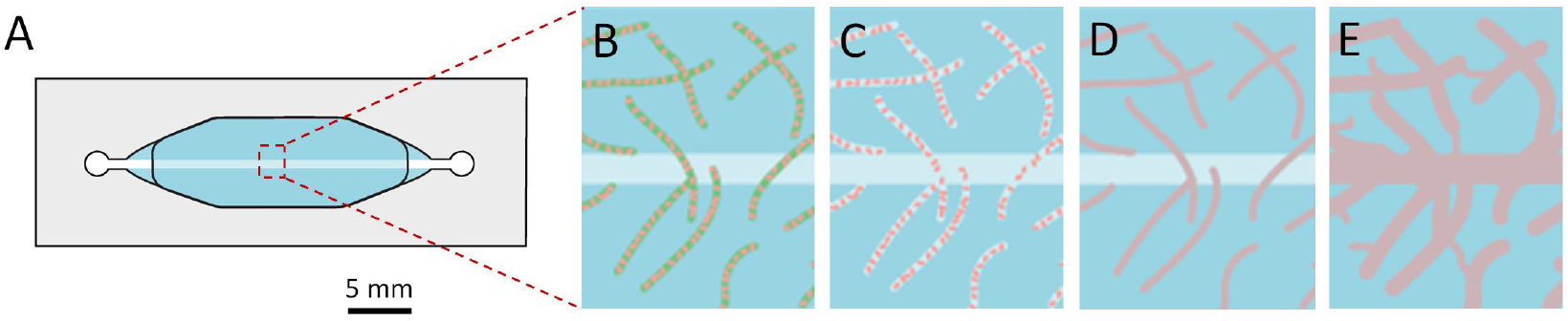
Schematic of microvessel network formation using sacrificial alginate threads. A) Millifluidic device with single channel. B) Alginate threads containing endothelial cells seeded in collagen-alginase solution. C) Alginate dissolved, releasing the cells into the newly-hollowed channels. D) Cells spread out to cover the channels left by the dissolved alginate. E) Capillaries sprout and connect to the main channel to create a microvessel network.

To embed the alginate threads in collagen, a frozen aliquot of GFP-HUVECs encapsulated within the alginate threads was quickly thawed in a 37°C bead bath. After aspirating the freezing media, the threads were transferred into the device chamber, which had a 25 gauge 2” length blunt tip needle (McMaster-Carr) inserted within the central channel (**Figure 3A)**. A solution of high concentration rat tail collagen was diluted to 5 mg/mL on ice and neutralized with 1 N NaOH, added 1 µL dropwise until neutral pH was demonstrated by the solution’s peach color. Alginase (Sigma Aldrich, St. Louis, MO) was added to the neutralized collagen to a final concentration of 0.05 UN/mL, including the volume of the alginate threads. The neutralized collagen/alginase solution was added to the device chamber over the alginate threads, suspending them in the collagen (**Figure 3B**). The final gel volume in the devices was approximately 250 µL.

To prepare control collagen gels containing randomly seeded cells, a solution of 1.5 million cells/mL was suspended in the neutralized collagen/alginase solution and then added to the device chamber.

In both conditions, the filled devices were placed on ice for 15 min to promote the formation of more optically transparent collagen gels that would be favorable for imaging. The devices containing gels were then incubated at 37°C for 30 min to fully solidify the collagen. In the condition of the millifluidic devices, before removing the needle, 200 µL warm media was added to the top chamber, a slab of PDMS was glued to cover the top, and the needle was back filled with media. When the needle was removed in this manner, the central channel remained intact. The needle hole in the PDMS on the side of the device was sealed with Loctite GO2 glue. In the condition of the well plate control, the samples were seeded and gelled as described above, but within a 24-well plate instead of a millifluidic device. In the well plate control, 2 mL of media was added on top of the gels after they were fully set.

### Sterile flow loop setup

To create the reservoir for the flow loop, a 3 mm hole was drilled into a 15 mL conical tube using a Dremel tool (Robert Bosch Tool Corporation, Racine, WI). Next, a 28 cm tube and a 22 cm microbore Tygon tube with 0.020” inner diameter and 0.060”outer diameter (Cole-Parmer, Vernon Hill, IL) were inserted into the cap with the ends offset 6 cm within the tube, and the tubes were secured with Loctite GO2 glue. The ends were offset to ensure that bubbles from the outlet were not recaptured at the inlet. After curing for 24 hours, the flow loop was assembled as shown in **Figure 2A**. To form the part of the flow loop that attaches to the device, a 14 gauge ½” blunt tip needle (McMaster-Carr) was inserted into the long Tygon tube attached to the reservoir and another into a loose 22 cm Tygon tube. To create the part of the flow loop that attaches to the pump, the Luer-lock ends of the needles were connected to a 11 cm long piece of 1/16” ID silicone tubing (Cole-Parmer) using 2x Luer-lock female-to-barb connectors (McMaster-Carr). The assembled loops were autoclaved before use.

### Application of flow

To connect the vascular network device to a flow system, a sterile flow loop was filled with warm media, then primed using a peristatic pump to bring the meniscus to the end of the loop closest to the reservoir, ensuring that perfusion into the device would begin upon pump activation (**Figure 2A**). The media formed a drop at the end of the tube that allowed it to be connected without introducing bubbles to the device. Bubbles elsewhere in the loop would be trapped by the reservoir. This end was pressure fit to the device and then glued. The other end was attached to the device in a similar way and then the pump was run to remove bubbles from the loop. The loop could be disassembled at the barb connectors to add holders for the peristaltic pump if desired. Flow was introduced to the devices at a rate of 710 µL/min. The media in the device reservoirs was exchanged with pre-warmed fresh media after 2 days, consistent with standard HUVEC culture. Flow was not introduced to the well plate controls.

### Imaging

Using a Nikon A1-Rsi confocal (Nikon Instruments Inc., Melville, NY) with a 10x objective, images were taken of 1.3 mm by 3.4 mm sections of the devices centered on the central channel approximately in the middle of the device, 2 cm from the outlet end. The closed loops were removed from the incubator for about 30 min at a time for this imaging, and then immediately returned to their static or flow conditions. This schedule allowed the imaging of the live cells in these devices and in well plates at 0, 1, 2, and 3 days post seeding. Large images were obtained using a Nikon Eclipse Ti2 (Nikon Instruments Inc., Melville, NY) with a 4x objective. Exposure time and laser power were kept consistent for all images in an experiment to simplify automated image analysis.

### Image analysis

Total cell volume was quantified using MATLAB (Mathworks, Inc.). A subset of images was used to manually set the fluorescence threshold for all images across experiments and timepoints. Cell count for the cell viability assays was performed using a CellProfiler pipeline and was validated with a manual count of cells that intersected a grid overlay.

### Statistical analysis

Quantitative data are shown as mean ± standard deviation. Statistical analysis was performed using GraphPad Prism 10.0.3 (San Diego, CA). A two-way analysis of variance (ANOVA) was used to compare the viability of HUVECs in threads versus in solution and to evaluate the effect of seeding method and perfusion over the culture duration using the millifluidic perfusion system. Tukey’s post-hoc testing was performed for comparisons between groups. Differences with a p-value of <0.05 met the threshold for significance.

## Experiment

### Cell viability test

Using the custom-designed, 3D printed coaxial extruder, HUVEC-laden alginate threads of approximately 330 ± 100 µm diameter and 4 ± 0.6 mm length were fabricated. Cell viability tests showed that cell-laden alginate threads could be frozen and thawed with minimal impact on cell viability, comparable to unencapsulated cells (**Figure 4A**). This was confirmed by two-way ANVOA of the variation between the freezing condition and encapsulation state. While freezing did significantly impact viability (p = 0.03), encapsulation did not (p = 0.54). The viability of encapsulated cells that had been frozen then thawed (76±10%) was not significantly different from the viability of unencapsulated cells that had been frozen then thawed (81±17%). The viability of the encapsulated cells and unencapsulated cells before freezing were 92±3% and 94±4%, respectively.

**Figure 4.**
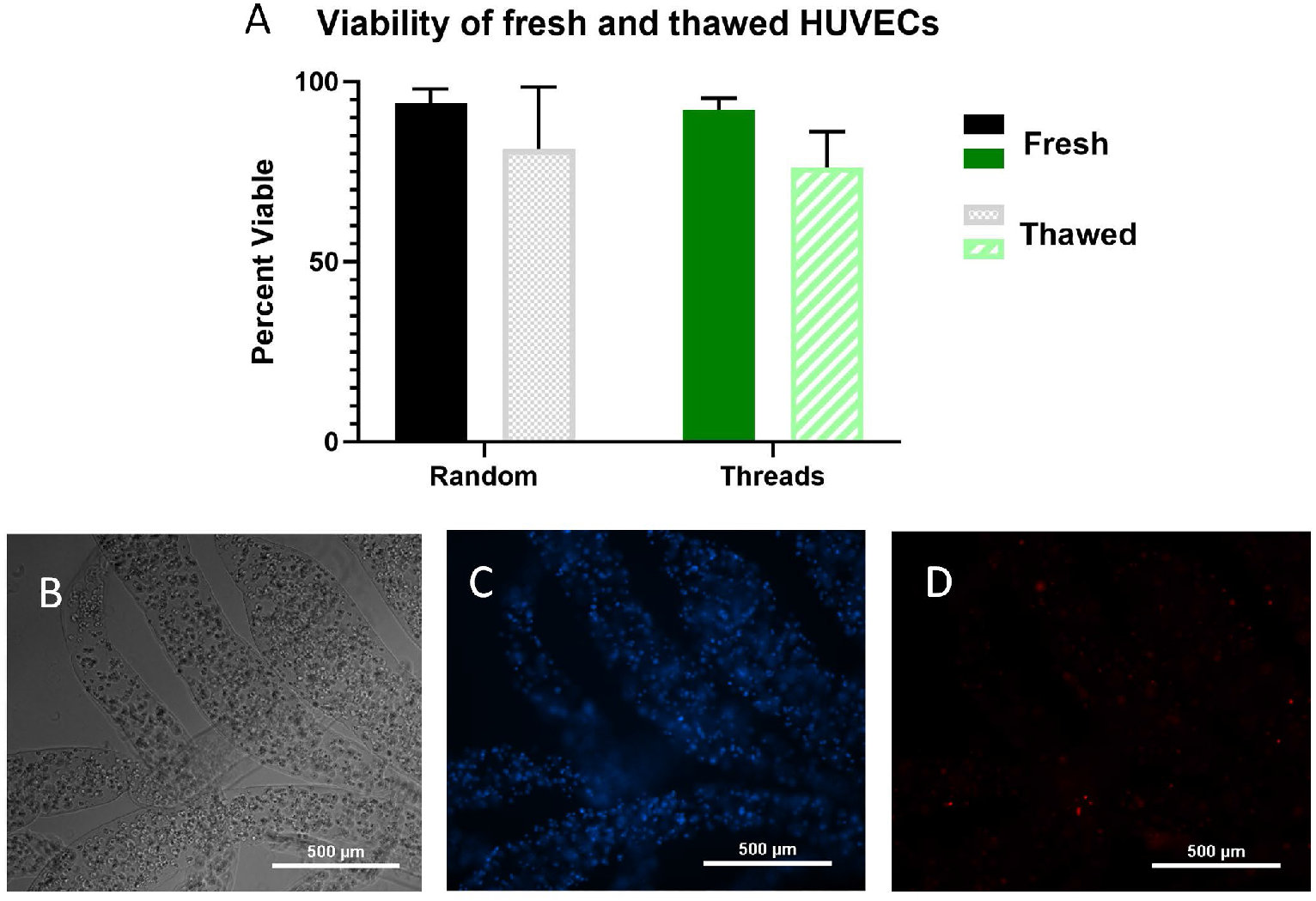
Viability of cells within alginate threads. A) Viability of HUVECs when passed, in threads immediately after extrusion, and after freezing then thawing. B) 10x image of HUVECs in alginate threads in phase. C) Calcein-AM live stain of HUVECS within alginate threads. D) Eth-D dead stain of HUVECS.

### Cell proliferation as a function of seeding method

After three days of culture within collagen gels, the cells within the well plates proliferated in both the random and alginate thread seeding conditions, nearly doubling in total cell volume (**Figure 5 and Supplemental Figure S1**) and were not significantly different between these two conditions (p = 0.94). Within the millifluidic devices under static conditions, cells in the alginate thread seeding condition proliferated after three days (p < 0.05), whereas the randomly seeded cells decreased (p < 0.01) (**Supplemental Figure S1**). Seeding with threads in the static condition yielded significantly increased cell proliferation compared to the randomly seeded samples under static conditions (p <0.05) (**Figure 5B**).

**Figure 5.**
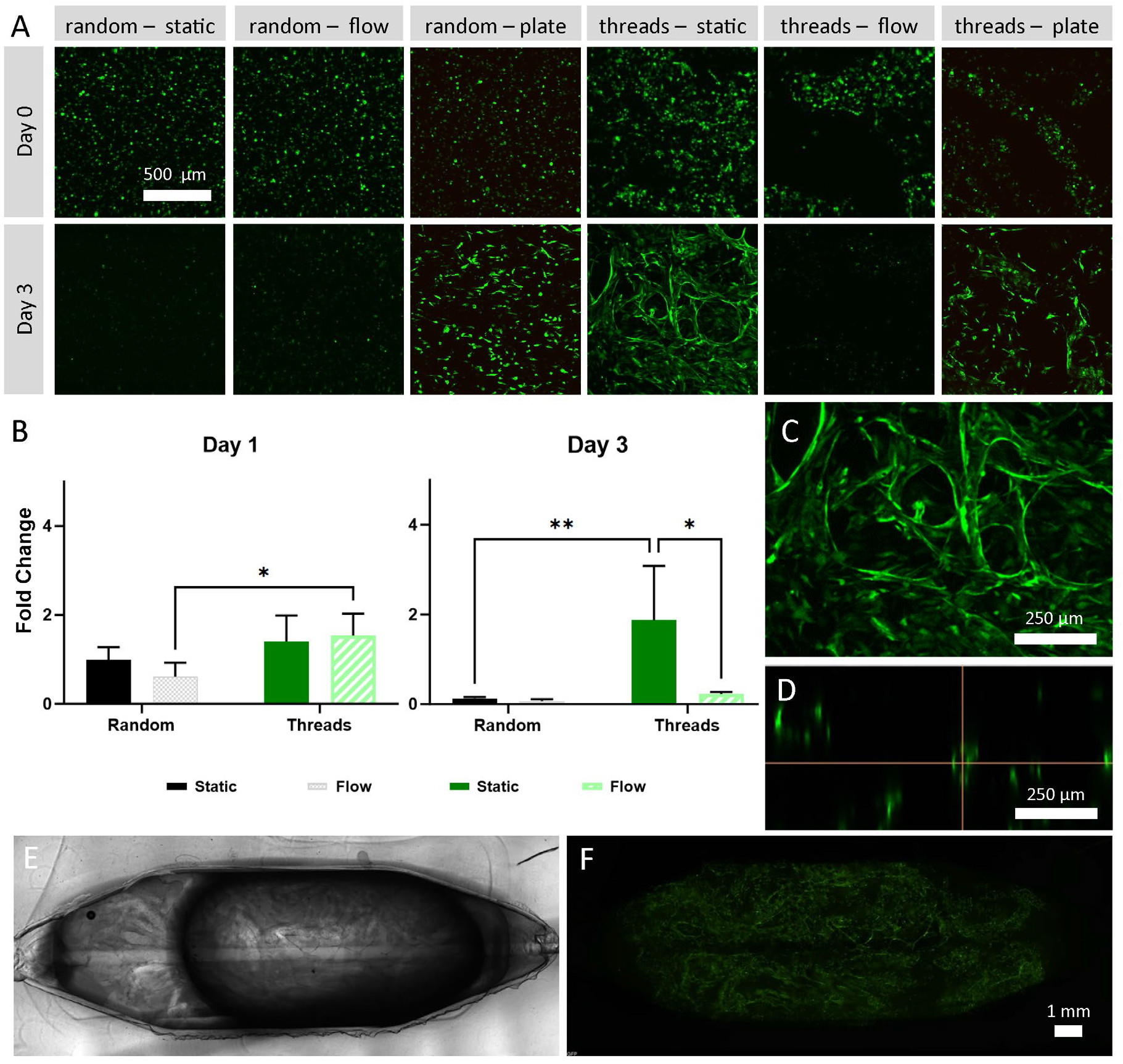
GFP-HUVECs encapsulated in collagen. A) Z-stack maximum intensity projections of GFP-HUVECs under conditions (top label), on Day 0 or Day 3 (left label). B) Fold change in GFP-HUVEC volume at Day 1 and Day 3. Each sample was normalized to its own Day 0 data. C) GFP-HUVECs seeded within threads after three days of culture magnified from lower right image in panel A. D) Z-stack cross section from C showing vessel lumen (diameter ∼30 µm). E) Phase image of millifluidic device culture chamber containing GFP-HUVECs seeded within threads and cultured for 3 days under static conditions. F) Corresponding GFP-HUVEC fluorescence from E. *p < 0.05; **p < 0.01.

In areas of high local cell density, new structures appeared to bridge between the channels formed by the threads (**Figure 5C**). Based on confocal imaging, these bridging structures appear to be capillaries of ∼ 30 µm diameter with a hollow lumen (**Figure 5C-D**). The functionality of these possible vessels could not be visualized with dye due to the blind-ended nature of the structures, so verification that these structures are hollow has not been completed. The device has an area of 1.28 cm^2^, with these capillary-like structures spanning the entire area (**Figure 5E-F**). These results indicate that alginate threads are useful for creating centimeter-scale networks of microvessel-like structures. In addition, the rapid formation of lumenized capillaries bridging the hollow alginate structures demonstrates that hierarchical network formation can be prompted by this patterning method.

### Cell proliferation as a function of flow

Under flow conditions, the cells in the alginate threads cultured in the device initially began to elongate and connect with neighboring cells, but by the third day, the fluorescent cell population decreased sharply (**Figure 5A-B**). Randomly seeded cells cultured within devices were nearly completely dead by their second day under flow conditions (**Figure 5A**). While there was a statistically significant difference between these two groups (p = 0.01), it is negligible because of the extensive cell death observed when the thread (p < 0.01) and random (p < 0.001) seeding conditions were subjected to flow (**Supplemental Figure S1**). At day 3 the difference between cell proliferation in the static and flow conditions in the threads seeding condition was significant (p < 0.0001). As early as day 2 in the random seeding condition, cell proliferation in both flow and static were significantly lower than cells randomly seeded in collagen gels in a well plate (p < 0.05). It is notable that the cells exposed to flow began to die off around the channel in the randomly seeded conditions, though the extent or large-scale geometry of this localized cell death was not captured by the image size. This flow-related cell death might be prevented by lowering the volume of media in the flow loop, changing the flow rate, or introducing flow at a later time point after the cells have fully recovered from being frozen.

## Discussion

Cells behave differently in 2D compared to 3D; therefore, it is important to develop vascularized 3D models of tissues. While other groups have used sacrificial structures to create single vessels^18^ or structures with beads^11^ or fibers^7^, the technique we present here results in a hierarchical microvascular structure and is easy to implement without expensive fabrication equipment. Other groups have also used coaxial flow to generate tubular cellular structures.^12,19^ However, our 3D printable coaxial extruder is easily transferrable to other labs and could be readily altered to create sacrificial structures of other dimensions.

In this report, we have described a compact, disposable flow loop including a built-in bubble trap and a reservoir for easy media changes. The inclusion of cell-laden alginate threads into the collagen gels under static conditions generated hierarchical microvascular networks, as would be necessary to support large, complex, 3D tissue models. The sacrificial threads can be frozen and thawed for use, making this system convenient and easy to implement in co-culture experiments. These results were in line with other encapsulation systems for cryopreservation.^20^ We showed that the viability of encapsulated cells were not significantly affected by the encapsulation process or freezing, compared with unencapsulated cells.

The cell seeding experiments within collagen gels demonstrated the utility of this method in generating microvascular networks. The collagen concentration and cell density have been optimized for capillary formation in previous work.^1^ We observed a lower degree of vascular morphogenesis than reported in the literature, however this can be explained by the shorter culture period and the recently unfrozen status of the cells. The difference in relative fluorescence observed between the starting conditions in random and threads samples (shown in **Supplemental Figure S2**) could be explained a difference in cell seeding conditions, but it is more likely that the alginate has a protective effect on the cells, acting as a barrier against mechanical forces, temperature, and pH changes.

The conditions within the well plate were sufficient to allow cell survival, whereas the device was only able to support the cells seeded within the sacrificial threads. The cells likely survived in this condition because the increased porosity of the collagen gel (∼30%) due to the voids formed by the sacrificial threads was sufficient to allow the diffusion of nutrients. Under flow, there was a sharp decrease in cell viability, possibly due to the washout of homeostatic factors or low-flow mediated apoptosis. Starting flow after the cells have completely recovered from freezing and grown to confluence, perhaps one week after seeding, may prevent this problem. All these results indicate that this technique for forming microvessels has utility in bioengineering for generating microvessel networks.

The networks resulting from this technique more closely resemble *in vitro* microvascular networks than the cellular structures created by a seemingly comparable technique that uses cell-covered gelatin beads,^11^ owing to the formation of distinct lumens instead of monolayers forming around voids between spheres and the potential to be able to adjust the diameter of the sacrificial structures. Similarly, the application of this technique also results in the approximate doubling of cells soon after seeding. Another notable work used a single sacrificial alginate vessel to create a channel within a millifluidic device.^18^ The technique presented here expands upon that concept to create a technique that is more suited toward generating large scale microvascular networks in a flexible format that can be used from frozen aliquots.

While developing this technique for successful formation of alginate threads used to make microvessel networks, there were challenges in the process that can be addressed in the future. Pipetting the threads with standard pipette tips can be challenging, which led to the design of the open-top design of the millifluidic device for the seeding process. Due to the high interstitial flow resistance of collagen at high concentrations, it was necessary to seal the open top of the device to prevent leakage and encourage flow through the microvessel networks. Improvements could be made to the design to encourage flow while facilitating the luminal and abluminal access necessary to perform permeability studies. Future studies, where microvessel networks are cultured to confluence, could include permeability studies and characterization of tight junctions. Our technique could also be adapted to co-culture vascular cells alongside perivascular cells, such as smooth muscle cells, which would further enhance the expression of tight junctions.

Though the threads were consistent in overall shape, the diameter of the threads varied along their length. Though this may not be a drawback when trying to form a network randomly, it would make it difficult to isolate the effects of vessel diameter on the network formation. The thickness of short alginate threads can be controlled by adjusting the speed of core extrusion, with slower speeds producing threads with greater diameter and faster speeds producing thinner threads, similar to adjusting the extrusion rate of materials used in electrospinning to control the diameter of the material. Shorter alginate threads could produce more uniform networks with lower connectivity. Similarly, longer alginate threads could produce more connected networks, but would be more difficult to distribute uniformly and repeatably. Optimizing the mix of different sized structures to create hierarchical microvessel networks with morphology similar to in-vivo networks is a topic worthy of further research.

In randomly oriented microvessel networks, the shear stress that the cells experience is not possible to control with precision and consistency, as parts of the network would experience turbulent flow and different flow rates as channel diameters varied. These challenges could be overcome by using fluorescent microbead tracking to measure the flow within different parts of a confluent microvessel network.

Lastly, since the threads are randomly oriented within the collagen gel, there is significant variability of the orientation and density of structures within the sample. Microvessel networks in vivo are hierarchical, but in vitro this non-homogeneity can make it challenging to reproduce experimental results. Reproducibility of the microvessel networks could be improved by using more advanced fabrication equipment and materials (e.g., 3D bio-printing, photocrosslinkable materials fabricated with high resolution DLP printers, traditional microfluidic and millifluidic fabrication) to create more uniform sacrificial structures. Additionally, to prevent settling of the structures, the composition of the threads and the matrix could be adjusted to be neutrally buoyant, or the collagen matrix replaced with a hydrogel material that crosslinks more rapidly. Scaling up the size of the devices or scaling down the size of the threads would be another way to improve the reproducibility of the microvessel networks.

In conclusion, here we present a method for generating sacrificial, alginate structures and embedding them within collagen to speed the formation of microvessel networks. The alginate threads were created using a 3D printed coaxial extruder and syringe pump; a simple method that is easily transferrable to other labs and flexible for others to tailor to their needs. The encapsulated cells retained viability within the alginate threads after encapsulation and after freezing. The flow loop and millifluidic device create a solid platform for studying in vitro tissues under recirculating flow. Experiments confirmed that the cells within alginate structures survive better than randomly seeded cells under static conditions with limited access to media, indicating that this technique could be useful in bioengineering applications with low levels of diffusion. The methods and tools presented here will contribute to the development and scale-up of lab-grown, vascularized tissues.

## Acknowledgements

The authors thank Linda Liu for assistance with device fabrication and Elysa Jui for statistical discussions.

## Author Contribution Statement

Conceptualization: SH, JC, and JG-A. Investigation: SH. Data analysis: SH and FA. Writing – original draft: SH. Writing – review & editing: SH, FA, and JG-A. Methodology: SH, JC, and JG-A. Supervision: JG-A. Funding acquisition: SH and JG-A. All authors contributed to the article and approved the final submitted version.

---

**Supplemental Figure S1.**
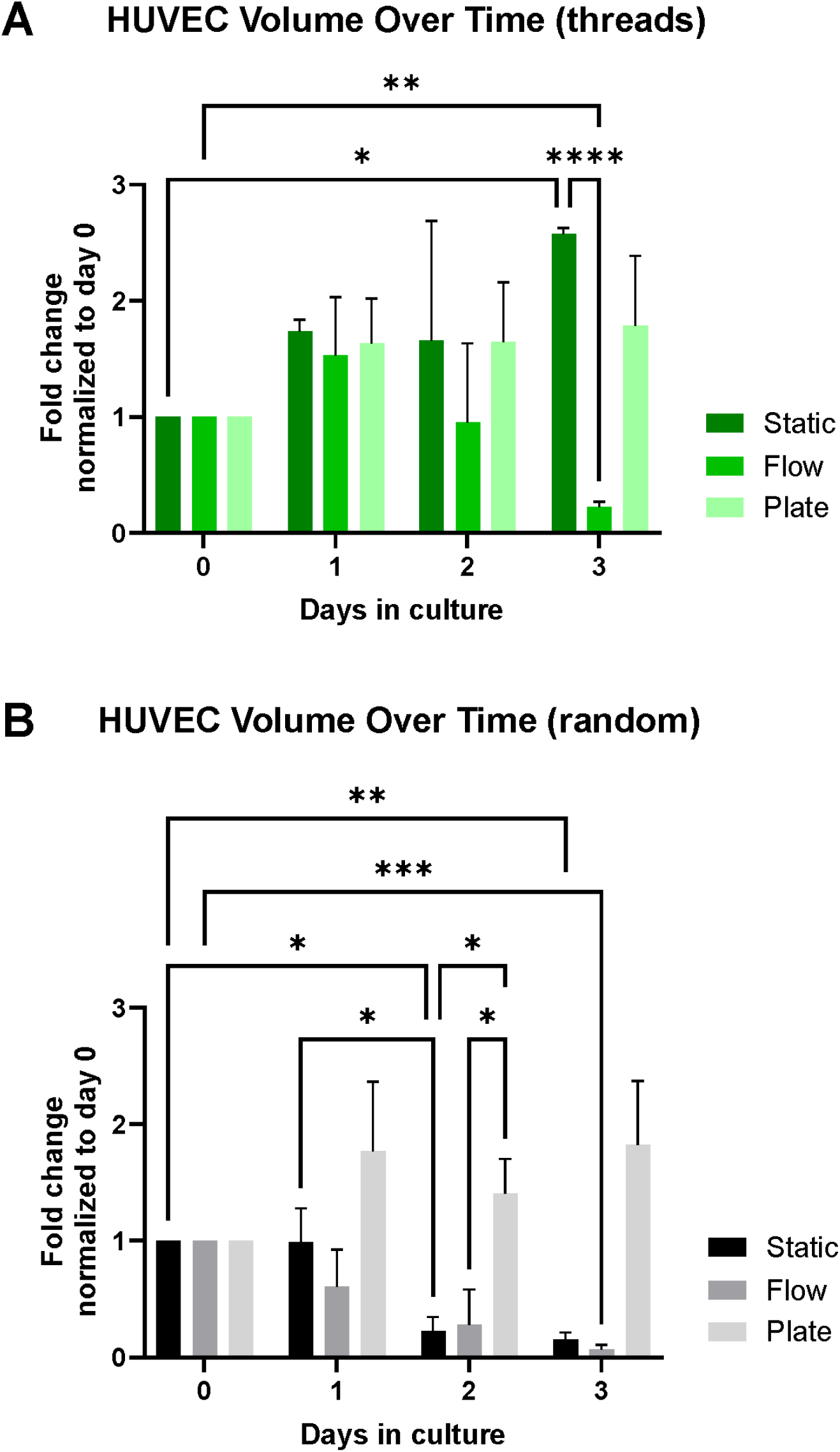
Total volume of GFP-HUVECS over time for static, flow, and well plate culture conditions in the alginate threads (A) and random (B) seeding conditions. Cell numbers for each day are expressed as fold change normalized to day 0 (statistical significance indicated: *p < 0.05, **p < 0.01, ***p < 0.001, ****p < 0.0001).

**Supplemental Figure S2.**
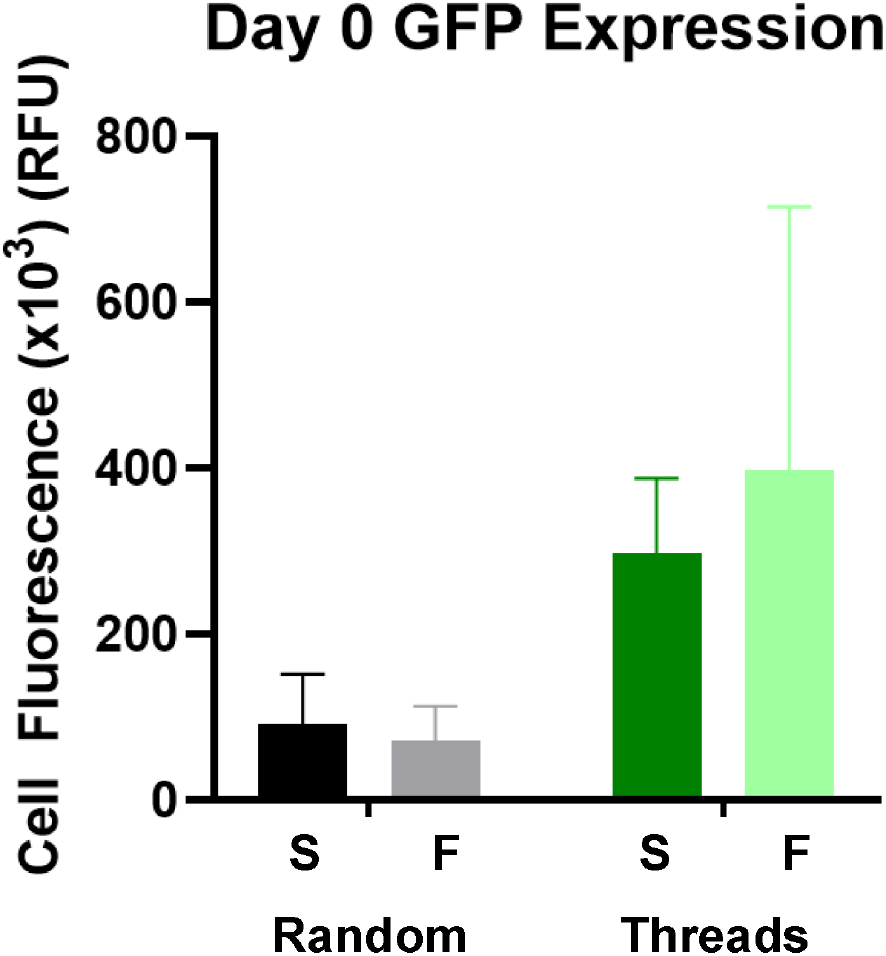
Relative fluorescent units (RFUx10^3^) GFP at day 0 for GFP-HUVECs seeded under random and threads conditions in preparation for culture under either static (S) or flow (F) conditions.

